# Heterogeneous origins of human sleep spindles in different cortical layers

**DOI:** 10.1101/221234

**Authors:** Donald J. Hagler, Istvan Ulbert, Lucia Wittner, Lorand Erőss, Joseph R. Madsen, Orrin Devinsky, Werner Doyle, Daniel Fabo, Sydney S. Cash, Eric Halgren

## Abstract

Sleep spindles are a cardinal feature in human slow wave sleep and may be important for memory consolidation. We studied the intracortical organization of spindles in humans by recording spontaneous sleep spindles from different cortical layers using linear microelectrode arrays. Two patterns of spindle generation were identified using visual inspection, and confirmed with factor analysis. Spindles were largest and most common in upper and middle channels, with limited involvement of deep channels. Many spindles were observed in only upper or only middle channels, but about half occurred in both. In spindles involving both middle and upper channels, the spindle envelope onset in middle channels led upper by ∼25-50ms on average. The phase relationship between spindle waves in upper and middle channels varied dynamically within spindle epochs, and across individuals. Current source density analysis demonstrated that upper and middle channel spindles were both generated by an excitatory supragranular current sink while an additional deep source was present for middle channel spindles only. Only middle channel spindles were accompanied by deep gamma activity. These results suggest that upper channel spindles are generated by supragranular pyramids, and middle channel by infragranular. Possibly, middle channel spindles are generated by core thalamocortical afferents, and upper channel by matrix. The concurrence of these patterns could reflect engagement of cortical circuits in the integration of more focal (core) and distributed (matrix) aspects of memory. These results demonstrate that at least two distinct intracortical systems generate human sleep spindles.

**Significance Statement:** Bursts of ∼14Hz oscillations, lasting about a second, have been recognized for over 80 years as cardinal features of mammalian sleep. Recent findings suggest that they play a key role in organizing cortical activity during memory consolidation. We used linear microelectrode arrays to study their intracortical organization in humans. We found that spindles could be divided into two types. One mainly engages upper layers of the cortex, which are considered to be specialized for associative activity. The other engages both upper and middle layers, including those devoted to sensory input. The interaction of these two spindle types may help organize the interaction of sensory and associative aspects of memory consolidation.

## Introduction

Spindle oscillations are a characteristic feature of slow wave sleep that were first described 80 years ago based on scalp electroencephalography (EEG) in humans (Loomis et al., 1935 ). They are bursts of ∼10-16 Hz activity lasting ∼0.5-2 seconds that occur during normal slow wave sleep (stages N2 and N3) (Luthi, 2013). Recent evidence supports a role for spindles in organizing replay of prior events as critical step in memory consolidation (Mednick et al., 2013, Rasch and Born, 2013). The basic neurophysiological mechanism of spindle generation involves intrinsic T and H currents in thalamic neurons and reciprocal connections between inhibitory cells in the thalamic reticular nucleus and bursting thalamocortical neurons (McCormick and Bal, 1997, Luthi, 2013). The thalamocortical cells project this rhythmic activity onto pyramidal cells, inducing currents which are then the proximal cause for spindles recorded in cortical local field potentials, EEG, and the magnetoencephalogram (MEG).

Spindles were originally considered to be global thalamocortical events. As measured with scalp EEG in humans(Dehghani et al., 2010, Dehghani et al., 2011a), or on the cortex of anesthetized cats(Contreras et al., 1996), spindles tend to be widespread and highly synchronous. Spindle synchronization is thought to arise from cortical feedback to the thalamic rhythm generators, as decortication leads to desynchronization of thalamic spindles (Contreras et al., 1996), and this mechanism is supported by computational modeling (Bonjean et al., 2012). Recent evidence, however, suggests that spindles are not homogeneous. Whereas scalp EEG spindles tend to be distributed and synchronous, simultaneous MEG measurements have found them to be asynchronous and focal (Dehghani et al., 2010, Dehghani et al., 2011a). This is consistent with intracranial recordings which find that most, but not all sleep spindles occur locally (Andrillon et al., 2011, Piantoni et al., 2017)

The thalamocortical projection is known to be organized into two systems, ‘core’ focal projections and diffuse ‘matrix’ projections (Jones, 1998, Zikopoulos and Barbas, 2007). , suggesting that local and global spindles may be mediated by core and matrix systems, respectively (Piantoni et al., 2016). Since core and matrix systems project differentially to different cortical layers (Zikopoulos and Barbas, 2007), this implies that different spindles would evoke different intracortical laminar profiles of local field potentials (LFP). Early work in cats suggested that the intracortical physiology of spindles were similar to with recruiting responses and thus reflected afferents from non-specific thalamic nuclei (Dempsey and Morison, 1941, Li et al., 1956). However, Spencer and Brookhart (1961) also found patterns resembling augmenting responses which are due to specific thalamocortical afferents, and suggested that both specific and nonspecific systems are engaged in spindle generation, often during the same spindle discharge. They recorded relative to a distant reference, which renders laminar localization of transmembrane currents problematic (Kajikawa and Schroeder, 2011). Kandel and Buzsaki (1997) reported that the main laminar pattern of spindles in unanesthetized rats resembled augmenting responses, but mentioned that other patterns were also present. Thus, these earlier studies, while suggestive, are limited in either their technical implementation or focus, and in any case were in different species which may have differently organized sleep spindles. We report here what appear to be the first laminar profiles of spontaneous sleep spindles, recorded using linear microelectrode arrays in epilepsy subjects, focusing on the question of multiple spindle generators.

## Methods

### Participants and data collection

5 subjects (15-42 years old; 3 female; Fig. 1) with long-standing pharmaco-resistant complex partial seizures participated after fully informed consent according to the Declaration of Helsinki guidelines as monitored by the local Institutional Review Boards. Subdural grid and strip electrode arrays were placed in order to confirm the hypothesized seizure focus, and locate epileptogenic tissue in relation to essential cortex, thus directing surgical treatment. The decision to implant, the electrode targets and the duration of implantation were made entirely on clinical grounds without reference to this research. Slow wave sleep was detected by the prevalence of generalized slow rhythms and spindles in cortical and scalp electrodes.

**Figure 1.**
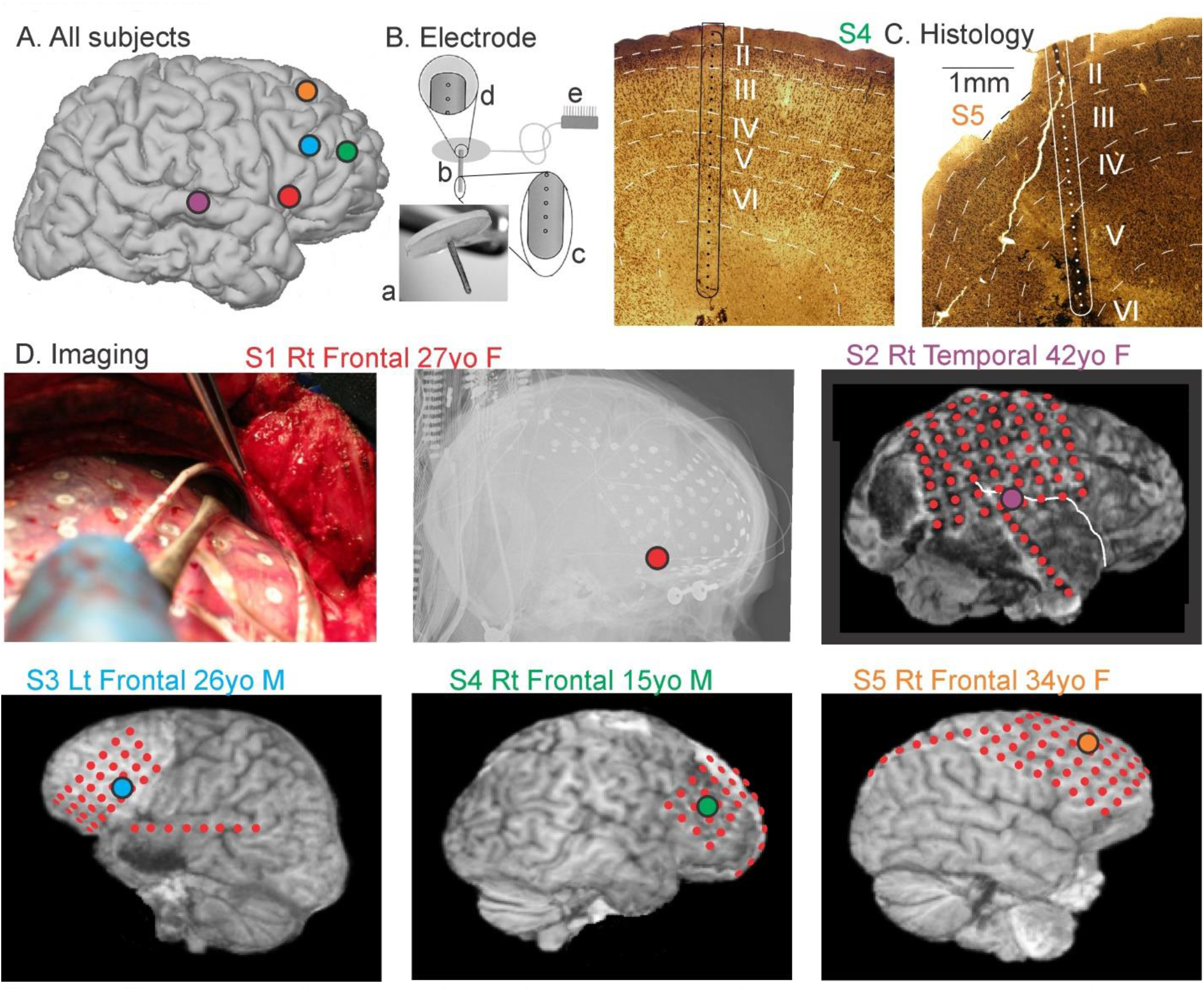
Electrode localization. A. Approximate locations of laminar probes are mapped on a standard right hemisphere. B. The laminar microarray: (a) photograph and (b) schematic of the entire array (3600μ long, 350μ diameter), and silastic flap placed on the cortical surface; (c) close-up of tip showing the five deepest contacts (each 40μ diameter, on 150μ centers); (d) close-up of the three most superficial contacts; (e) connector to amplifiers. C. Histology of the implant sites in two subjects, stained with NeuN. D. Imaging of electrode locations, showing placement of the laminar probe under direct visualization in subject 1 together with post-implant x-rays, and post implant MRIs in subjects 2 through 5. The hemisphere and lobe implanted, age and gender of the five subjects are also shown.

### Electrodes and Localization

A laminar microarray containing 24 90%Pt-10%Ir contacts, each 40μ, in diameter, at 150μ, center-to-center spacing (Ulbert et al., 2001), was placed under the grid in cortex that had been previously identified as probably epileptogenic, in the center of the likely surgical target in the frontal or temporal association cortex (Fig. 1A,B,D). Gyral localization was based on direct visualization during surgery in all subjects, supplemented with transmission X-ray (Subject 1), or structural MRI with electrodes in place (Subjects 2-5), and by post-implantation CT superimposed on pre-implantation MRI (Subjects 3-5), using a previously described procedure (Yang et al., 2012).

Localization with respect to cortical lamina was based on surgical procedure and electrode design, confirmed by histology in two subjects (Fig. 1C). The neurosurgeon placed the electrode under direct visualization, perpendicular to the cortical surface. The electrode is designed with a thin silastic flap which adheres by surface tension to the pial surface, and the electrode is kept in place because it is underneath the clinical grid and dura. Since the micro-contacts are known distances from the underside of the silastic flap, this constrains their distance from the pial surface. Specifically, the first contact was centered ∼150μm below the pial surface, and the 24^th^ contact at ∼3600μ below the pial surface. While this does not allow absolute localization of the contacts with respect to cortical lamina, it does permit their correspondence to be estimated from previous measurements of laminar width in human cortex (Hutsler et al., 2005).

In two of the five subjects it was possible to remove *en bloc* the cortex immediately surrounding the microelectrode array at the therapeutic cortectomy. Histological analysis of the microelectrode tracks was performed as described earlier (Csercsa et al., 2010). Briefly, tissue blocks were fixed with 4% paraformaldehyde, 0.1% glutaraldehyde and 0.2% picric acid in 0.1M phosphate buffer (PB). Sixty μm thick sections were cut with a Vibratome, thoroughly washed with PB, immersed in 30% sucrose for 1 -2 days, and then frozen three times over liquid nitrogen. Endogenous peroxidase was blocked by 1% H2O2 in PB for 10 minutes. Non-specific immunostaining was blocked by 5% milk powder and 2% bovine serum albumin. Mouse monoclonal antibody against the neuronal marker NeuN (1:3000, Chemicon, Temecula, CA, USA) was used for two days. For visualization of immunopositive elements, biotinylated antimouse immunoglobulin G (1:300, Vector) was applied, followed by avidin-biotinylated horseradish peroxidase complex (ABC; 1:300, Vector). The immunoperoxidase reaction was developed by 3,30-diaminobenzidine tetrahydrochloride (DAB; Sigma), as a chromogen. Sections were then osmicated (0.25% OsO4 in PB) dehydrated in ethanol, and mounted in Durcupan (ACM, Fluka).

Including all patients with laminar electrodes, histological localization of laminar electrode contacts was available in 9 subjects, of which two (subjects 4 and 5) are included in this paper. Surgical constraints did not allow *en bloc* resection of the implantation site in the other subjects. The 9 subjects with histology include one that was (intentionally) placed in a tuber, and one which was not perpendicular to the surface (subject 5 of the current paper). Omitting these two subjects, the mean and standard deviation of the center contact in each layer were: layer 1, contact 1.00±.00; layer 2, contact 3.43±.53; layer 3, contact 7.00±1.00; layer 4, contact 10.14±1.21; layer 5, contact 14.00±2.00; and layer 6, contact 19.14±3.39. The insertion point of the electrode in subject 5 was displaced from the crown of the gyrus and thus was at an angle to the cortical layers. Consequently, the central contacts of the layers in this subject were: layer 1, contact 2; layer 2, contact 5; layer 3, contact 10; layer 4, contact 14; layer 5, contact 21; and layer 6, not sampled. The electrode tracks in all of the other 8 subjects were perpendicular to the layers, and the surgeon in subjects 1-4 of the current paper attempted to place the electrodes in the gyral crown. Given the consistency of the overall histological results, and the lack of individual histological localization in three of the five subjects, we elected to analyze the data with respect to contact, and to use the minimal labels of ‘upper,’ ‘middle’ and ‘lower’ rather than the 6 canonical layers. Channels 1-4, labelled ‘upper’ would include layer 1 and parts of layer 2; Channels 10-14, labelled ‘middle’ would include layer 4 and parts of surrounding layers. This conservative approach is based on the known contact-layer relationship imposed by the physical constraints of the implantation and electrode design, and confirmed by histological analysis when available. Data are thus presented as averages by channel across subjects (figures 3, 4, 7, 8), as well as individual subject data to permit variability to be evaluated (figures 5, 6, 6, 8). This approach is further validated by the similarity of the laminar LFPg and CSD profiles across subjects (figure 5).

### Recordings and Preprocessing

Local field potential gradient (LFPg) recordings were made from 23 pairs of successive contacts. After wideband (DC-10,000 Hz) pre-amplification (gain 10×, CMRR 90db, input impedance 10^12^ ohms), the signal was split into LFPg (filtered at 0.2-500 Hz, gain 1,000x, digitized at 2,000 Hz, 16 bit) and action potentials (filtered at 200-5,000 Hz, gain 1,000x, digitized at 20,000 Hz, 12 bit), and stored continuously. Differential recording permitted adequate SNR on active wards due to high CMRR of radiated noise common to adjacent leads (Ulbert et al., 2001). Furthermore, modeling studies demonstrate that the LFPg between contacts at 150μm centers assures that activity is a reliable probe of locally generated activity, unlike referential LFP (Kajikawa and Schroeder, 2011). Current Source Density (CSD) is also reliably local, but is a derived measure with multiple assumptions and more susceptible to noise, gain equalization and other instrumentaion issues. Consequently, both LFPg and CSD data are displayed.Data were collected in an Epilepsy Monitoring Unit during naturally-occurring nocturnal sleep. Epochs were chosen for analysis when the patient was behaviorally asleep and widespread slow waves (delta activity) were present in the electrocorticogram, identifying slow wave sleep, i.e., stages N2 and N3. SWS was chosen rather than focusing on N2 or N3 because sleep spindles occur in both N2 and N3 (Andrillon et al., 2011, Mak-McCully et al., 2017, Piantoni et al., 2017), and to increase comparability to rodent studies where the N2/N3 distinction is not clear. An average of 50 min sleep (standard deviation = 21, range = 16-67) was analyzed per subject.

The recordings often exhibit various electrical artifacts, for example caused by the participant changing head position. Being epilepsy subjects, abnormal spiking activity is also possible. Furthermore, defective electrode contacts result in unusable recordings from two adjacent differential amplifier channels. To handle such artifacts, laminar electrode data were preprocessed to filter line noise, exclude high amplitude spikes, and replace “bad” channels through interpolation and slight spatial smoothing. To suppress line noise, a notch filter was applied (zero phase shift, frequency domain), with a stop frequency of 50 or 60 Hz (depending on acquisition site) and a transition band equal to 30% of the stop frequency. This means that for a stop frequency of 50 Hz, oscillations between 50 and 65 Hz were gradually attenuated, until being completely suppressed above 65 Hz. “Bad” channels, which typically exhibit frequent, high amplitude artifacts, usually occurring with opposite polarity in two adjacent channels, were identified visually for each laminar recording (i.e. the time series data collected with a laminar electrode array for a single subject). There were between 2 and 10 of these bad channels in each recording (mean = 5.4, SD = 3.0; S1: [17, 18]; S2: [13, 14, 22, 23]; S3: [1, 3, 4, 12, 13, 15, 16, 19, 20, 23]; S4: [13, 14, 18, 19, 23]; S5: [7, 10, 16, 17, 19, 20]). Time windows of high amplitude fluctuations were excluded using a manually determined masking threshold. For each laminar recording, time series data were visually reviewed (excluding “bad” channels) and a threshold value was chosen to be greater than the normal fluctuations from the mean but less than the sharp, high amplitude deviations from the mean. This threshold was applied to create a binary (0 or 1) time series mask. Time points within 2 s of threshold crossings were set to 0, as were time windows shorter than 5 s in duration between potential artefactual periods in order to eliminate periods with repeated abnormal activity. Data for bad channels were replaced by the weighted average of neighboring, good channels, with weights that decrease with increasing spacing between channels according to an exponential decay function (decay constant = 0.1 channel spaces). One-dimensional spatial smoothing was then applied across channels (Gaussian sigma = 0.64 channel spaces, equivalent to FWHM = 1.5). This slight spatial smoothing was applied to ensure gradual and continuous variation across channels, which is commonly done to suppress false sources and sinks due to slight signal fluctuations. An average of 7.5% of the analyzed sleep period (standard deviation = 6.4%, range = 2.1-16.4%) was rejected per subject.

### Spindle detection

Spindles were detected using custom methods developed to accommodate the observed characteristics of spindles in laminar potential gradient recordings. The spindle detection algorithm, described in detail below, was based on the standard criteria of sustained power in the spindle band (Andrillon et al., 2011). Sensitivity was increased by relaxing amplitude and duration criteria, while selectivity was maintained by adding rejection criteria from adjacent bands (Mak-McCully et al., 2017), and validating with visual inspection (Figure 1). Time courses of spindle-band amplitude were calculated using a zero phase shift, frequency domain, bandpass filter (10-16 Hz), with transition bands equal to 30% of the cut-off frequencies. Absolute values of the filtered data were smoothed across time by convolving with a tapered cosine (Tukey (Harris, 1978) window (width = 300 ms, ratio of cosine taper to constant region = 0.5). To account for between-channel differences in spindle-band amplitude, channel-specific, median values of spindle band amplitudes were subtracted from each channel. Offset spindle-band amplitudes were then normalized by dividing by a robust estimator of the standard deviation (rSD: the median absolute deviation divided by 0.6745). To preserve relative differences in spindle amplitudes between channels, rSD values were averaged across channels and then applied as a uniform normalization factor.

Potential spindles were identified independently for each channel by finding peaks in the normalized spindle-band envelope time courses with amplitudes greater than 1. Spindle onset and offset were estimated by finding the nearest time points before and after the peak of the spindle-band envelope with amplitudes less than 1. Possible spindles were excluded if the estimated spindle durations were less than 200 ms. Epochs containing a possible spindle in one or more channel were then identified, and the overall spindle epoch onset and offset were estimated as the earliest onset and the latest offset across channels.

To avoid filtering artifacts such as ringing, and to adequately account for the reported variation in spindle frequency, a relatively broad band pass filter was used to identify spindle epochs. Consequently, this method is sensitive to high amplitude, transient increases in power at frequencies that are slightly lower or higher than the spindle range of 10 to 16 Hz. For this reason, we excluded events with relatively high amplitude in lower and higher frequency bands. To exclude such epochs, low-band (4-8 Hz) and high-band (18-25 Hz) time courses were calculated as described above for the spindle-band, except with different band-pass filters. Epochs were rejected if they contained low-or high-band amplitudes greater than 5 (relative to the rSD of low-or high-band amplitude) between the supposed spindle onset and offset in any of the channels containing a supposed spindle for that epoch. This strategy also rejected theta bursts, which have a frequency profile which often extend into spindle range, and can be seen to occur prior to downstates in Figure 2, where spindles occur after downstates.

**Figure 2.**
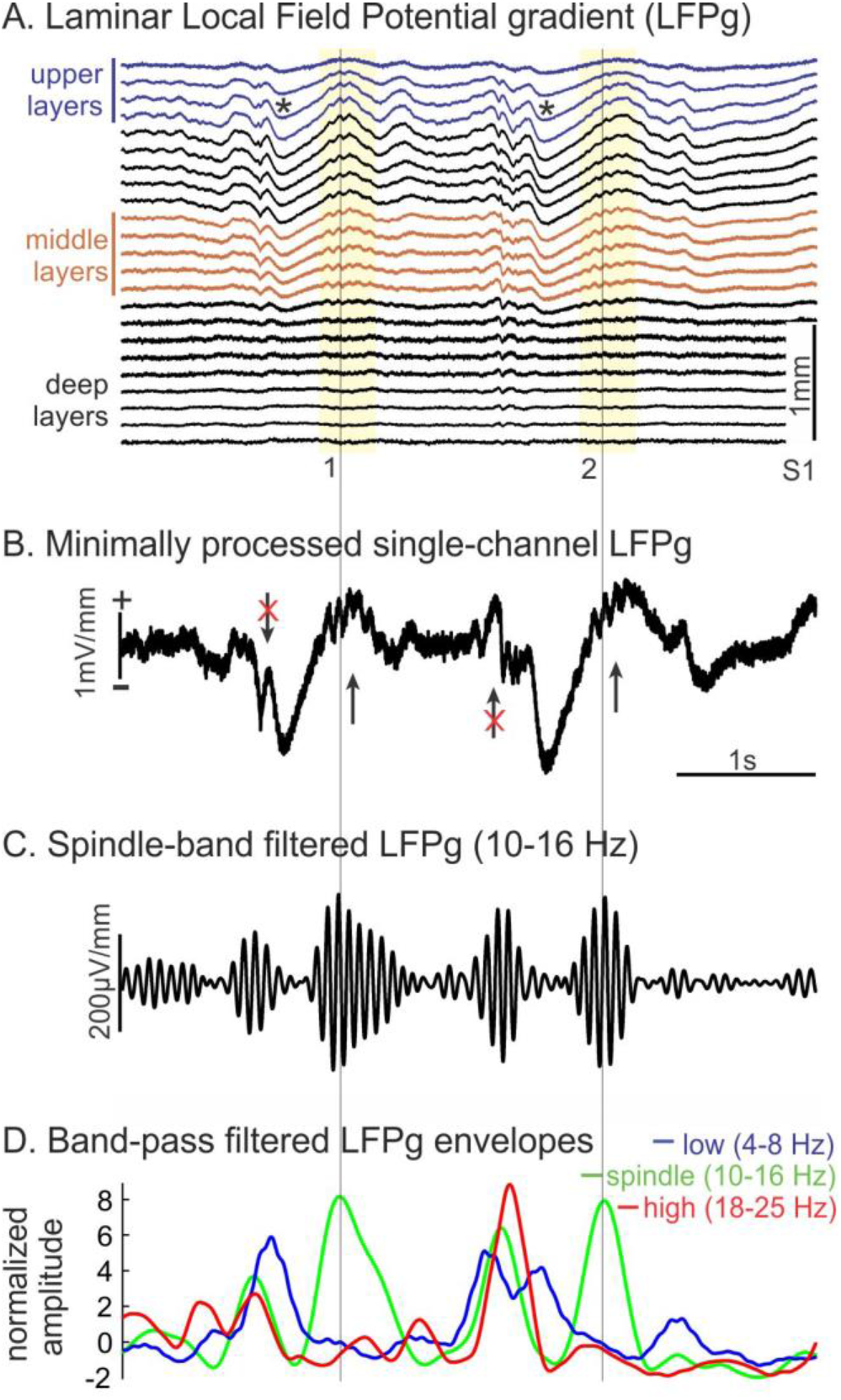
Spindle detection and patient characteristics., A. Example of laminar recordings of two spindles (1, 2) following downstates (*). The first spindle is clear in both upper channels (blue traces) and middle channels (brown traces), whereas the second is only clear in middle channels. Data were minimally preprocessed:60 Hz notch filter, interpolation to replace bad channels and slight spatial smoothing. B. Spindles detected in one middle channel are marked with arrows. Rejected spindles, with excessive low or high frequency oscillations are marked with arrows with red X’s through them. C. Band-pass filtered data in spindle band (10-16 Hz) from middle layer channel. D. Normalized amplitude envelopes of low-(4-8 Hz), spindle-, and high-band (18-25 Hz) oscillations.

Spindle identification inevitably results in false positives and false negatives, especially near the detection threshold. False negatives are a particular concern in evaluating spindle spread between layers, where they could result in an underestimation of the degree of intracolumnar spread. In order to determine whether characteristics of spindles were robust to the choice of detection threshold, we initially used a relatively low amplitude threshold in order to detect spindles in as many channels as possible, and then repeated our analyses including only spindles with overall amplitudes above the median. The overall amplitude of a given spindle epoch was defined in this context as the maximum across channels of the peak spindle-band amplitude. Spindle co-occurrence rates for both analyses are reported, but our primary analyses focused on the “strong” spindles with amplitudes greater than or equal to the median amplitude. An average density of 13.1 strong spindles per minute (standard deviation = 2.0, range = 10.5-15.8) were detected across subjects.

### Current Source Density (CSD)

Population trans-synaptic current flows were estimated using Current Source Density (CSD) analysis (Nicholson and Freeman, 1975). As preprocessing specific to CSD calculations, additional one-dimensional smoothing was applied across channels (sigma = 0.85, FWHM = 2). CSD time courses were then calculated from the smoothed LFPg data by applying a regularized inverse (SNR = 20) matrix operator calculating the second spatial derivative (Pettersen et al., 2006). The equivalent current dipole (ECD) was calculated from the estimated CSD by essentially calculating a distance-from-center weighted average of the CSD, implemented using trapezoidal integration. Note that one-dimensional CSD assumes that the extracellular tissue conductance does not change over the sampled trajectory, and that the transmembrane currents are symmetrical around that trajectory.

### Phase amplitude coupling

Spindle epochs were sorted into two groups based on detection in upper channels but not middle, or in middle channels but not upper. For each channel group, a reference channel was selected based on highest fraction of spindles detected for a given recording. We define the fraction of spindles detected as the number of spindles detected in a given channel relative to the total number of spindles detected in any channel. Centered on each spindle (with spindle epoch center determined by the average of estimated epoch onset and offset), 4000 ms epochs were filtered for three different frequency bands (spindle: 10-16 Hz; low gamma: 25-50 Hz; high gamma: 70-170 Hz) using a zero phase shift, frequency domain, band-pass filter, with transition bands equal to 30% of the cut-off frequencies. Amplitude and phase were obtained from the Hilbert transformation. To avoid edge artifacts, and to limit analysis to the stronger part of the spindle, the epoch was then reduced to -400 to 400 ms relative to the center of each spindle epoch. Spindle phase and gamma amplitudes were sampled from a single CSD-derived channel corresponding to the peak of the current sink. Spindle phases for each epoch were sorted into 32 bins (10 degrees) and low and high gamma amplitudes were averaged across time points within each phase bin.

To visualize this phase-amplitude relationship (Fig. 4), spindle time courses and low and high gamma amplitude were averaged across epochs for each channel, time-locked to the largest spindle-band peak in the channel with the highest fraction of spindles detected, for epochs that were either in upper channels only or middle channels only, and then averaged across subjects (n=5). Gamma amplitudes were transformed into z-scores before averaging across subjects, using the average and standard error of gamma power across all epochs between -400 and 400 ms of spindle center.

### Phase differences between channels

Spindle epochs were selected based on detection in both upper channels and middle channels. One reference channel was selected for each channel group, as described above for phase amplitude coupling analysis. The average phase difference for each subject was calculated for the selected upper channel relative to the middle channel. Each spindle epoch was re-centered in time to the highest amplitude spindle peak in the selected middle channel for that epoch. Instantaneous phase was derived from Hilbert transforms of band-pass filtered (10-16 Hz) waveforms. Phase differences were calculated at each time point by subtracting the instantaneous phase value of the middle channel from the instantaneous phase value of the upper channel. Phase differences between -200 ms and +200 ms were averaged using the circular mean (circ_mean) function of the CircStat MATLAB Toolbox (Berens, 2009). These average phase differences were then averaged across epochs, again using a circular mean. Because the polarity of LFPg measurements in individual channels depend on that channel's position relative to sources and sinks, it is unclear how to correctly interpret phase differences approaching a half cycle, since this could reflect polarity inversion, rather than a consistent phase lag in one channel relative to another. For this reason, for phase differences that were less than -0.25 cycles or greater than +0.25 cycles, an additional 0.5 cycles were added or subtracted, respectively, from the calculated phase difference (for subjects 2, 3, and 5).

### Experimental design and statistical analysis

Epochs containing laminar recordings of spontaneously occurring spindles were detected in slow wave sleep in 5 subjects. In addition to describing these unique recordings, with standard descriptive statistics, we performed the following statistical analysis to determine if gamma amplitude is modulated by spindle phase. The Modulation Index (MI) was calculated as described by Tort et al., 2010, from the phase-amplitude-coupling (gamma amplitude for each phase bin). Bootstrap resampling across epochs with 2000 iterations was used to estimate 95% confidence intervals for MI (Efron, 1981). Although these measures are ideally zero under the null hypothesis, spurious, non-zero, phase-amplitude relationships can occur. To estimate the rate at which this occurs, we generated a highly conservative, upper-bound null hypothesis value using the upper 95^th^ percentile value of MI calculated with phase information scrambled across epochs. On each of 100 iterations, gamma amplitudes of each epoch were assigned to the spindle phases of a randomly chosen epoch, based on random permutations of the epoch order. MI was calculated for each of these permutations, forming a distribution to represent the null hypothesis. The 95^th^ percentile value of this distribution was estimated from the mean plus twice the standard error. The bootstrap confidence intervals were then compared to this upper-bound null value in the calculation of p-values. Separate analyses were performed for spindles which were limited to either upper or middle channels.

When spindles occurred in both middle and upper channels, then a one-sample t-test was performed with one value from each subject to compare the latency of spindle onset (as described above). These latencies were determined in each co-occurring spindle individually, the signed differences obtained, and averaged across all such spindles for that subject, and the resulting mean was used for the t-test. The same procedure was used to compare the latency to peak in upper vs middle channels. Paired t-tests with two values from each subject were used to compare the proportion of spindles detected in upper channels that also occurred in middle channels, to the proportion of spindles detected in middle channels that occurred in upper.

Principal component analysis (PCA) was performed on the data for each subject using singular value decomposition (SVD), as implemented in the runpca function included in the EEGlab toolbox (Swartz Center for Computational Neuroscience, La Jolla, CA). Input data were spindle-band (10-16 Hz) filtered laminar time series data, concatenated across spindle epochs, with time points before and after the estimated spindle onsets and offsets excluded. Spatiotemporal patterns of spindle components were visualized by calculating average spindle time courses for each of the first two principal components. For these calculations, the Eigenvectors and Eigenvalues returned by the PCA analysis for each of the two components were individually applied to the spindle-band filtered data to obtain sensor-space time courses for each component. Spindles were averaged across epochs, time locked to the largest spindle peak in each epoch, and then averaged across subjects. The Eigenvector returned by PCA for a given component is a set of weights for each laminar channel. The sign of the Eigenvector being arbitrary and variable across subjects, we flipped the sign if necessary to make the mean of the channel weights for a given Eigenvector to be positive, allowing for a meaningful group average.

## Results

Laminar electrode recordings collected from five human epilepsy subjects were included in the current study (see **Methods**: *Data collection).* Laminar recording sites were located in either frontal or temporal lobes. Epochs containing a spindle in at least one of the 23 laminar channels were identified based on the amplitude envelope of band-pass filtered activity between 10-16 Hz (see **Methods**: *Spindle detection,* Fig. 2). The maximum spindle amplitude across channels was calculated for each putative spindle epoch we detected. To reduce the possibility of including non-spindle events in our analyses, epochs with maximum amplitudes less than the median maximum amplitude were excluded from our primary analyses. Spindles were visible in the raw traces, often following K-complexes (Cash et al., 2009) or downstates (Csercsa et al., 2010). Typically, distinct LFPg profiles were noted for downstates *versus* spindles, but this was not quantitatively examined (Fig 2A).

The laminar profile of spindle amplitudes varied considerably across spindle epochs. Some spindles were restricted to a few adjacent channels in either the upper range (e.g. between channel 1 and 4) or the middle range (e.g. 10-14), although many spindles were detected simultaneously in both upper and middle channels (Fig. 3A). Averaged across epochs, spindle amplitude was greatest in middle channels (average normalized spindle amplitude: 3.5 ± 0.2 SEM, n=5), slightly weaker in upper channels (2.3 ± 0.1 SEM), and weaker still in deep channels (1.2 ± 0.4 SEM) (Fig. 3B). Laminar variation in the likelihood of detecting a spindle was closely related to spindle amplitude, with most frequent detection in middle channels (average fraction of spindles detected: 0.72 ± 0.04 SEM), somewhat reduced frequency of detection in upper channels (0.52 ± 0.03 SEM), and further reduced fraction of spindles detected in deep channels (0.25 ± 0.10 SEM).

**Figure 3.**
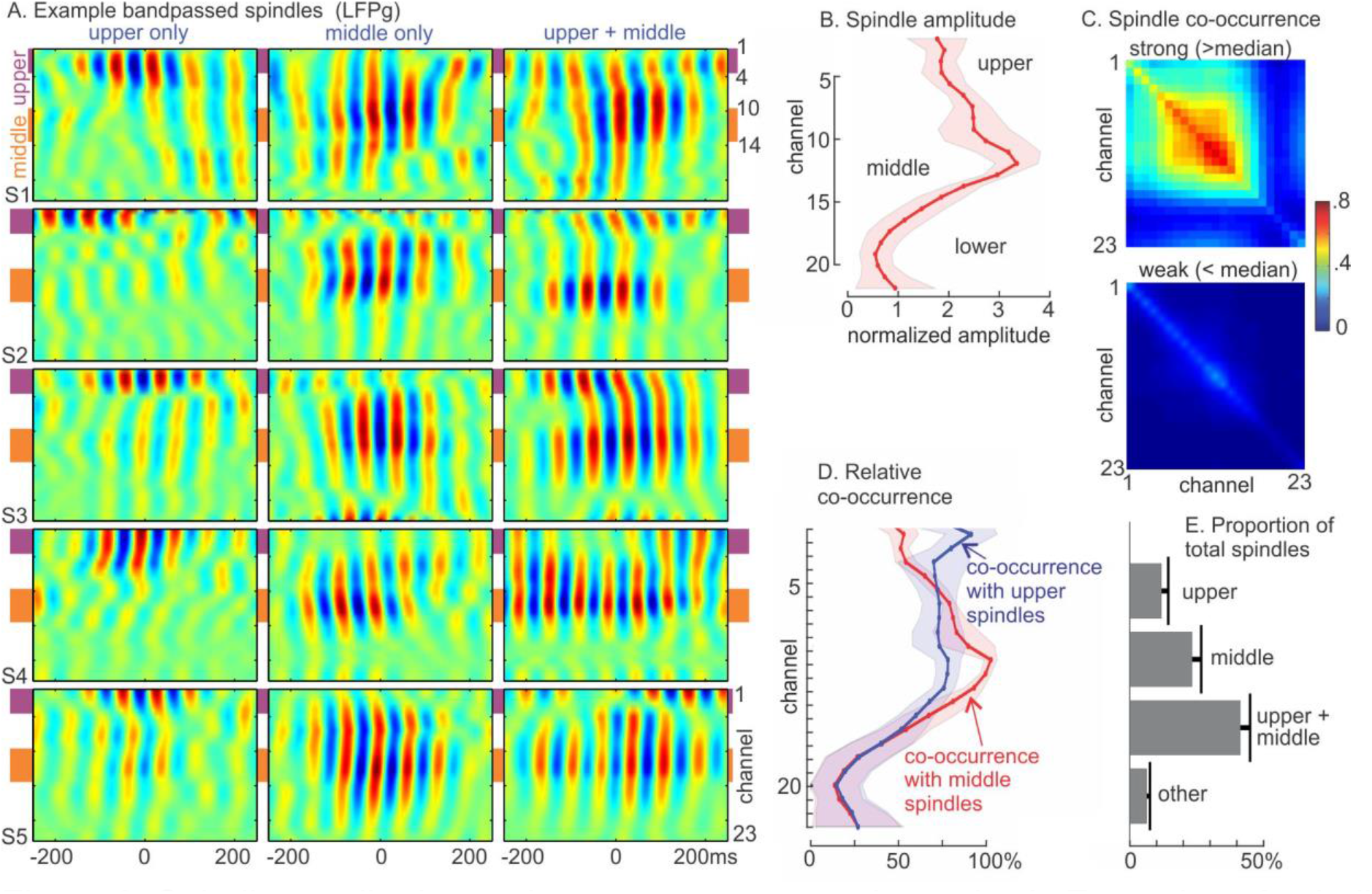
Spindle amplitudes and occurrence across channels. A. Examples of spindle activity (band-pass filtered, 10-16 Hz) detected in upper channels only, middle channels only, or both upper and middle channels. B. Group average (n=5) of peak normalized spindle-band amplitude averaged across all epochs for each channel. Shaded region indicates 95% confidence interval (1.96 × SEM). C. On top, spindle co-occurrence matrix for “strong” spindles, with maximum amplitude greater than the median. On bottom, co-occurrence matrix for “weak” spindles. Displayed values represent the number of “strong” or “weak” spindles detected in a given channel or pair of channels divided by the total number of “strong” or “weak” spindles detected in any channel for a given subject. Thus, the diagonal indicates the proportion of total spindles that included that particular channel. The plot suggests that strong spindles usually involved multiple middle channels, whereas weak spindles tended to occur in a single channel. D. Relative co-occurrence rates for epochs containing a spindle in either upper or middle channels. E. Group average of the fraction of spindles detected in upper channels only, middle channels only, upper and middle channels, or other channels.

Most spindles were detected simultaneously in multiple channels, often with substantial spatial separation between the channels (Fig. 3C). For the higher amplitude, “strong” spindles, laminar recordings exhibited distinctive patterns of spindle co-occurrence - i.e. the rate at which a spindle was detected in a pair of channels in the same epoch. For epochs excluded based on lower than median amplitudes, co-occurrence was much lower, such that “weak” spindles were often detected in only one channel at a time. For spindles detected in upper channels (selected for each laminar recording based on highest fraction of spindles detected among channels 1 to 4), spindles were found to co-occur in middle channels more than half the time (0.76 ± 0.04 SEM; Fig. 3D). For spindles detected in middle channels (channels 10 to 14), spindles co-occurred in upper channels slightly less often (0.55 ± 0.03 SEM; difference from upper-detected channels p = 0.015, paired t-test; Fig. 3D). Spindles occurred in both upper and middle channels about half of the time, but sometimes occurred in only upper or only middle channels, demonstrating a degree of independence of occurrence (Fig. 3E).

We characterized the current sources and sinks associated with spindles detected in either upper or middle channels. The transmembrane currents generating the spontaneous spindle oscillations were estimated from the local field potential gradient (LFPg) using current source density (CSD) analysis (see **Methods**: *Current Source Density).* We used phase-amplitude coupling between spindle oscillations and gamma band amplitude to identify the spindle phase related to active current sinks, reflecting the increase in firing related to excitatory synaptic input (see **Methods**: *Phase amplitude coupling).* The laminar distribution of sources and sinks associated with spindles detected in either upper or middle channels, averaged across subjects, is shown in Fig. 4. To visualize phase-amplitude relationships, spindles were averaged across epochs, time-locked to the largest spindle peak in a reference channel chosen from the upper or middle channels based on having the highest fraction of spindles detected in that range.

**Figure 4.**
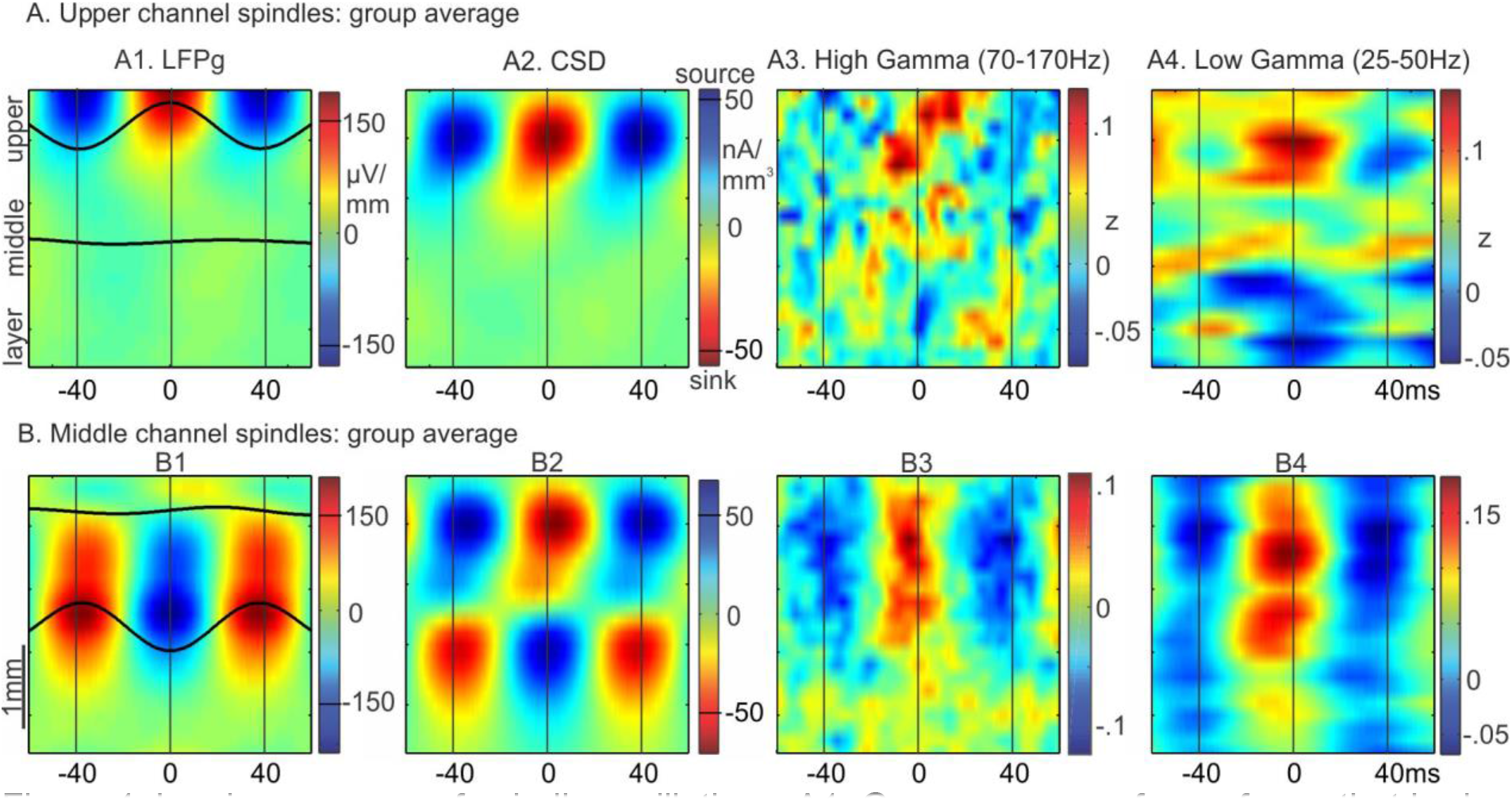
Laminar sources of spindle oscillations. A1. Group average of waveforms that had been averaged across epochs and individual peaks (band-pass filtered, 10-16 Hz), time locked to spindle peaks in the upper channel (channels 1-4) with the highest spindle rate for each subject. Only above-median upper channel spindle epochs were included. Waveforms are superimposed on a time-amplitude plot of LFPg in the same epochs. Time-amplitude plots from these epochs are also shown for: A2-CSD; A3-high gamma (70-170 Hz) amplitude; and A4-low gamma (25-50 Hz) amplitude. B. The same plots are shown, except in this case averages were made for middle channel spindle epochs time locked to the middle channel (channels 10-14) with the highest spindle rate for each subject.

For spindles detected in upper channels only, a positive peak in the LFPg corresponded to a current sink (negative CSD) in upper layers (e.g., 1 or 2/3; Fig. 4B1). For spindles detected in middle channels only, a negative peak in the LFPg corresponded to a current sink in upper layers and a current source in deep layers (Fig. 4B2). For each, around the time of those spindle peaks, there was an increase in both high gamma power (70 - 170 Hz; Fig. 4A3 and 4B3) and low gamma power (25-50 Hz; Fig. 4A4 and 4B4). For the upper channel spindles, there is the suggestion of a decrease in high gamma in lower channels at the time of the upper channel increase (Fig. A3, A4), as has been found in rat medial prefrontal cortex (Peyrache et al., 2011). The coupling between spindle phase and high gamma amplitude, summarized by MI, was significantly different from the null (p<0.05, bootstrap resampling) in four of the five subjects for upper channel spindles and for three of five for middle channel spindles (Table 1).LFPg and CSD patterns for upper and middle channel spindles varied across subjects, but showed overall consistency in the source/sink patterns (Fig. 5). For two subjects, there was an additional current source in the uppermost channels (Fig. 5A, S1 and S3), suggestive of a return current in layer I. Its apparent absence in the other three subjects may reflect variation in the thickness of this layer. Overall, these results suggest that excitatory input to upper layers can results in at least two different patterns of current sinks and sources, manifesting as positive LFPg peaks in upper channels or negative LFPg peaks in middle channels.

**Table 1.** Modulation of high gamma by spindle phase: Statistical results of comparison to the null distribution using bootstrap resampling. The normalized modulation index (MI) is a ratio that compares the calculated MI to the reference distribution calculated under the null hypothesis of no phase relationship. p-values were derived from bootstrap sampling of MI in comparison to the null for each of five recordings obtained from five different patients. The number of epochs varied across recordings, limited by factors such as the duration of good quality recording and the respective incidence of spindles in either the upper or middle channels but not both.

| Subject | Upper channel spindles |  |  | Middle channel spindles |  |  |
| --- | --- | --- | --- | --- | --- | --- |
|  | *number*<br>*of*<br>*epochs* | *normalized*<br>*modulation*<br>*index* | *p-value* | *number*<br>*of*<br>*epochs* | *normalized*<br>*modulation*<br>*index* | *p-value* |
| **1** | 81 | 2.03 | **0.0075** | 175 | 0.93 | 0.22 |
| **2** | 134 | 5.34 | **0.0005** | 93 | 3.26 | **0.001** |
| **3** | 85 | 3.85 | **0.001** | 246 | 4.46 | **0.0005** |
| **4** | 19 | 0.37 | 0.78 | 80 | 1.94 | 0.051 |
| **5** | 46 | 1.67 | **0.041** | 139 | 3.32 | **0.0015** |

**Figure 5.**
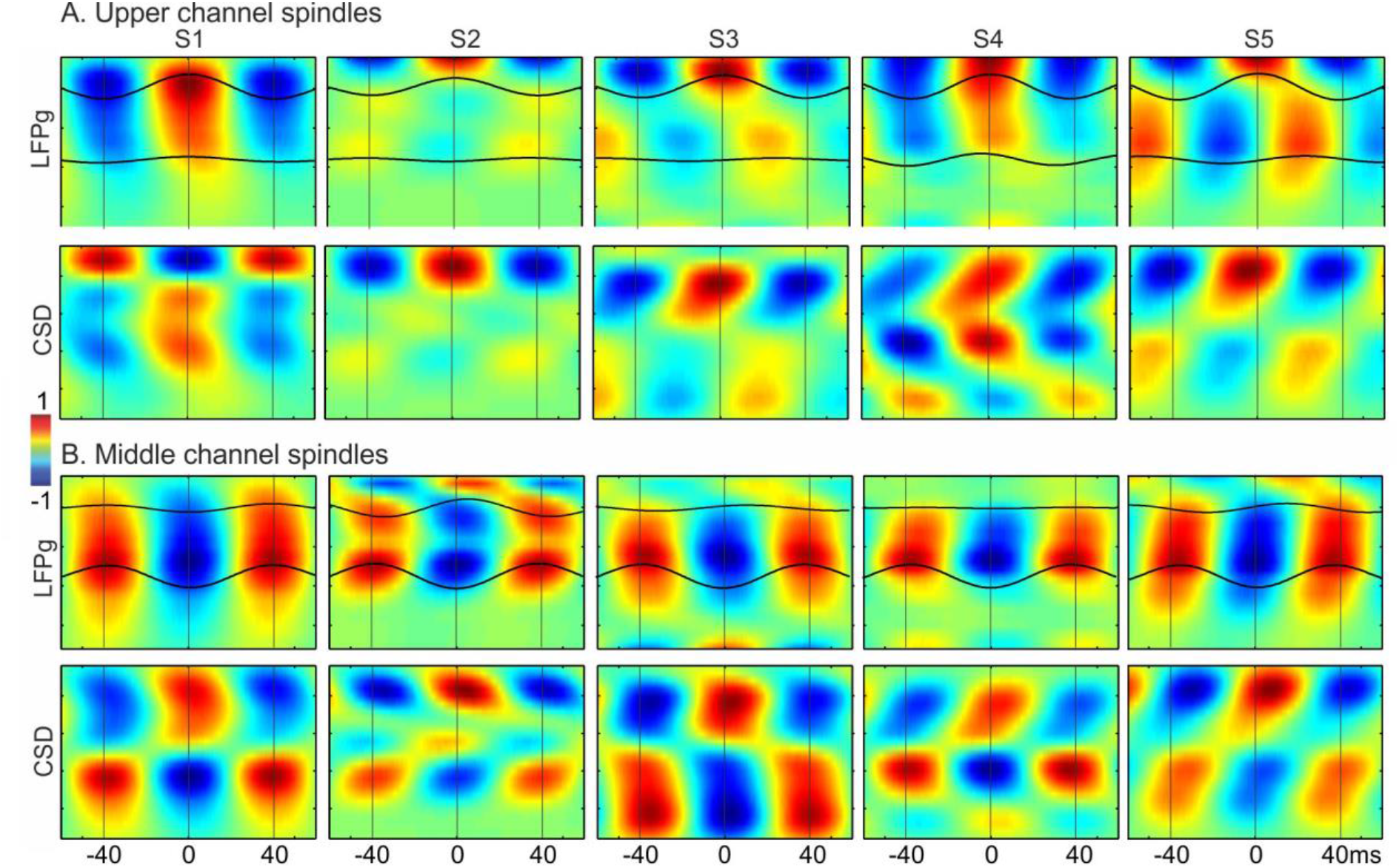
Laminar sources of spindle oscillations in individual subjects. A. Individual subject (arranged left to right) waveforms averaged across epochs and individual peaks (band-pass filtered, 10-16 Hz), time locked to spindle peaks in the upper channel (channels 1-4) with the highest spindle rate for each subject, with LFPg on top and CSD on bottom (both normalized to respective maximum values within each subject). B. Individual subject average waveforms time locked to the middle channel (channels 10-14) with the highest spindle rate for each subject.

We analyzed the relative timing and phase relationship between upper and middle channels for spindles detected simultaneously in both. One upper and one middle channel were selected for each subject based on maximum fraction of spindles detected in each range of channels (i.e. 1 to 4 and 10 to 14). Latencies of the onset of spindle epochs and peak of spindle-band activity were estimated from the smoothed, spindle-band amplitude envelope (see **Methods**: *Spindle detection).* Between-channel onset and peak latency differences were averaged over epochs (mean: 243 epochs, SD: 113, min: 69, max: 361). Because onset detection based on fixed thresholds could be biased by differences in the amplitudes of the signals, the threshold chosen for each channel and epoch was half of peak spindle amplitude. Based on the Hilbert transform analytic amplitude (i.e., amplitude envelope), spindle onset and peak latency tended to be delayed in upper channels (onset latency difference: 29.4 ms ± 10.1 SEM, p = 0.028, one sample t-test; peak latency difference: 18.2 ms ± 5.4 SEM, p>0.05, one sample t-test) (Fig. 6A).

**Figure 6.**
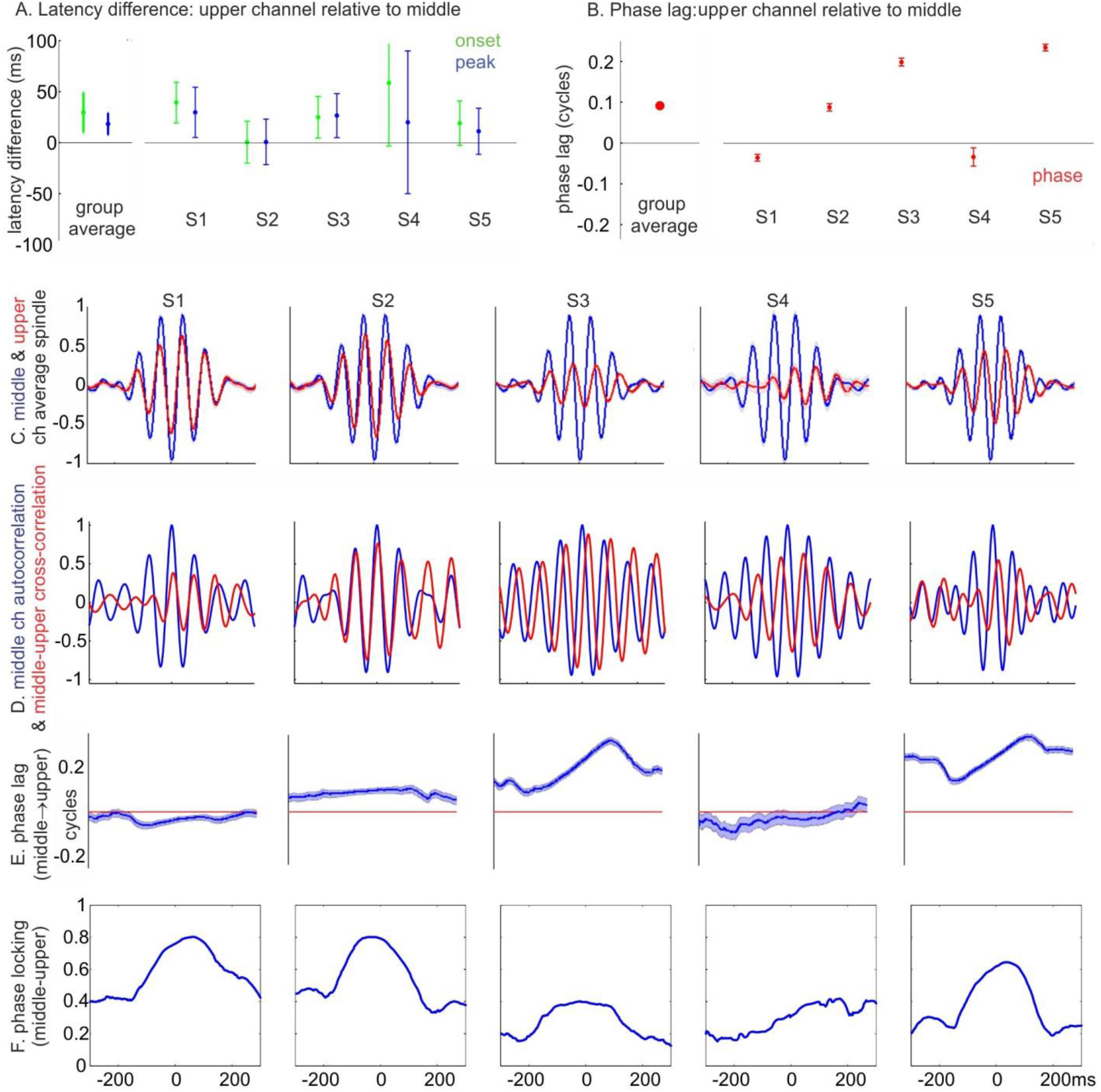
Spindle timing differences between channels. A. Latency differences between upper and middle channels for spindle epoch onset and peak of spindle-band amplitude envelope, averaged across spindle epochs. On left, group average. On right, individual subjects. Error bars indicate 95% confidence interval (1.96 × SEM). B. Phase lag of upper channels relative to middle channels (selected for each subject based on spindle rate). C. Normalized spindle waveforms averaged across epochs, time-locked to the middle channel (channels 10-14) with the highest spindle rate for each subject (arranged left to right). Middle channel in blue, upper channel in red. D. Cross-correlation of selected upper and middle channels for each subject, averaged across epochs (auto-correlation for middle channel in blue, cross-correlation between upper and middle channel in red). E. Phase lag averaged across epochs. F. Phase locking. In panels B-E, the upper layer spindles in subjects 2, 3, and 5 were inverted prior to plotting because their responses were >.25 cycle displaced from the middle layer spindles, suggesting phase-wrapping (see *Methods*).

Phase differences of individual waves between upper and middle channels were derived from Hillbert transforms of spindle band (10-16 Hz) filtered epochs, averaged across time points between -200 and 200 ms relative to spindle peak, and averaged across epochs (see **Methods**: *Phase differences between channels).* The group average phase lag was not significantly different from zero (p>0.05, one-sample test for the mean angle, using circ_mtest from CircStat Toolbox). The individual subject phase lags were significant for four of the five subjects (p<0.05, one-sample test for the mean angle, using circ_mtest from CircStat Toolbox) (Fig. 6B). In order to examine this relationship, the average of upper layer spindles relative to middle layer (Fig. 6C), their cross-correlation (Fig. 6D), and the time course of their phase-lag (Fig. 6E) and phase-locking (Fig. 6F), were calculated and plotted for each subject across all spindles. Although the patterns varied for unknown reasons, in some subjects, a consistent delay is observed which varies over the course of the spindle. Varying phase lags could be indicative of differences in oscillation frequency. Note that, in order to correct for presumed phase-wrapping, we effectively inverted polarity by adding or subtracting a half cycle of phase if the phase delay was more than one quarter cycle from zero in Fig. 6B-E. Despite this rationale, we acknowledge that flipping polarity in this way is somewhat arbitrary, particularly if the phase difference is only slightly more than one quarter cycle. For this reason, and because of inter-subject variability, it is difficult to draw conclusions about the directionality of the phase difference between upper and middle channels.

From the group average Fourier spectra, there appears to be a slight decrease in oscillation frequency for upper channel spindles (Fig. 7). However, differences between upper and middle channels in the peak spindle frequency during upper only or middle only spindles, respectively, were not significant (upper mean: 13.9 Hz, SD: 1.0, middle mean: 13.0 Hz, SD: 1.5, p>0.05, paired t-test).

**Figure 7.**
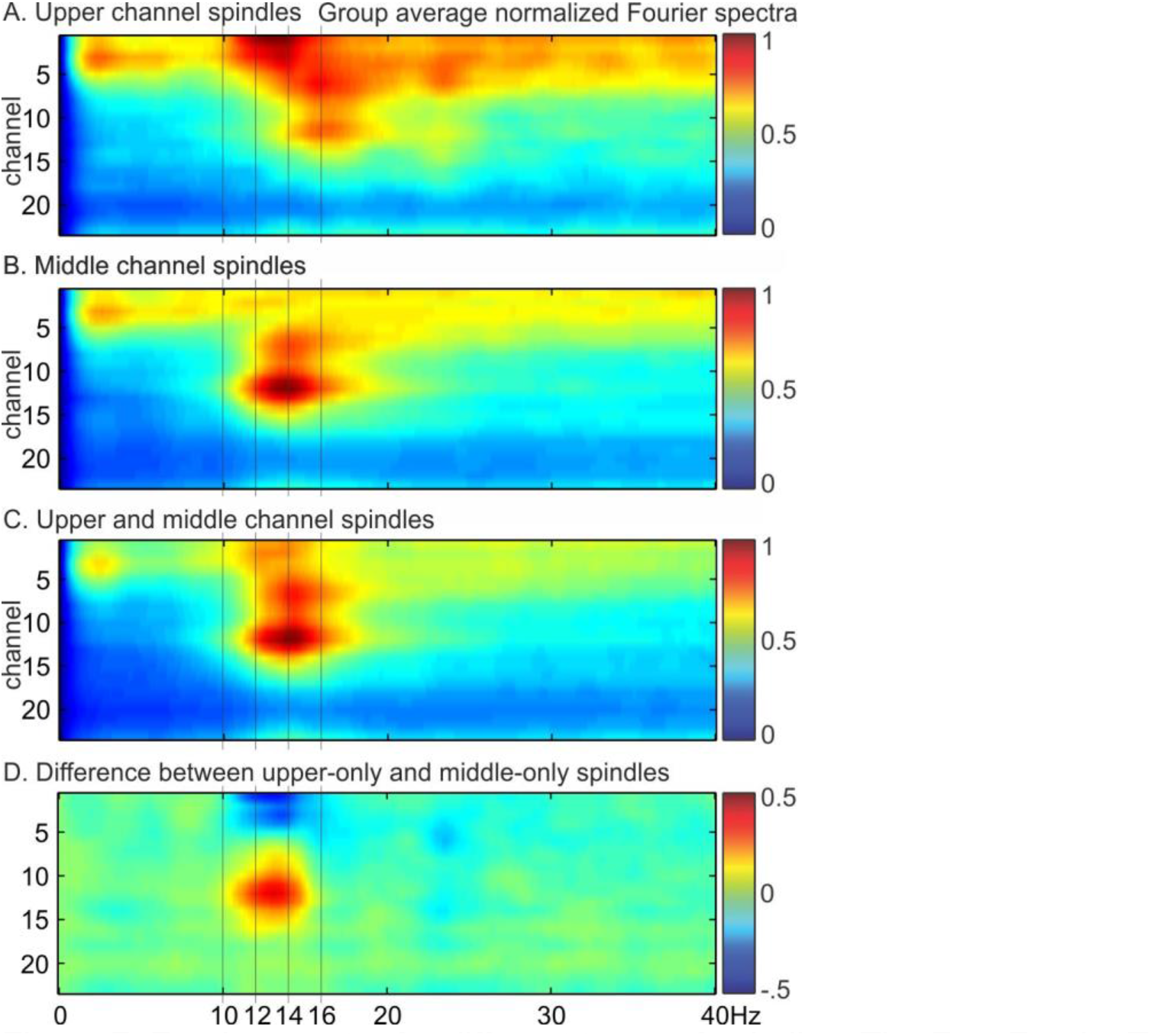
Fourier spectra for different types of spindles. The Fast Fourier Transform (FFT) was used to calculate spectral power during spindle epochs for each channel, averaged across epochs, normalized to the maximum power, and then averaged across subjects (n=5). A. Spindles detected in upper channels but not middle channels. B. Spindles detected in middle channels but not upper channels. C. Spindles detected in both upper and middle channels. D. Difference spectra between spindles detected in middle- or upper-channels only.

To analyze the spatiotemporal variation both across and within spindle epochs with a data-driven approach, we used Principal Components Analysis (PCA) to transform spindle-band activity from the 23 laminar channels into temporally uncorrelated components (Fig. 8). Spindle band filtered data from all spindle epochs (excluding “weak” spindles), delimited by estimated spindle onsets and offsets, were concatenated into one large matrix of channels by time points. The shape of the scree plot, showing the explained variance for each component, suggests that a two factor solution is appropriate (Bryant and Yarnold, 1995). The first two principal components (PCs) accounted for about half of the explained variance (Fig. 8A). The patterns of LFPg and CSD for the first PC (Fig. 8B) was very similar to that of spindles time-locked to middle channels (Fig. 4B), and the second PC was very similar to the upper channel spindles (Fig. 4A). These patterns are based on the component loadings, which were strong and highly similar across subjects for principal components 1 and 2, but weak and variable for later components (Fig. 8D). However, it is possible that with a larger dataset, one or more consistent patterns could be identified across multiple subjects. The degree of involvement of the first two PCs varied independently across epochs, and within epochs in which both PCs had strong involvement, the different PCs combined in various ways that accounted for the large variation in the dynamic laminar distribution of individual spindle epochs (Fig 9C).

**Figure 8.**
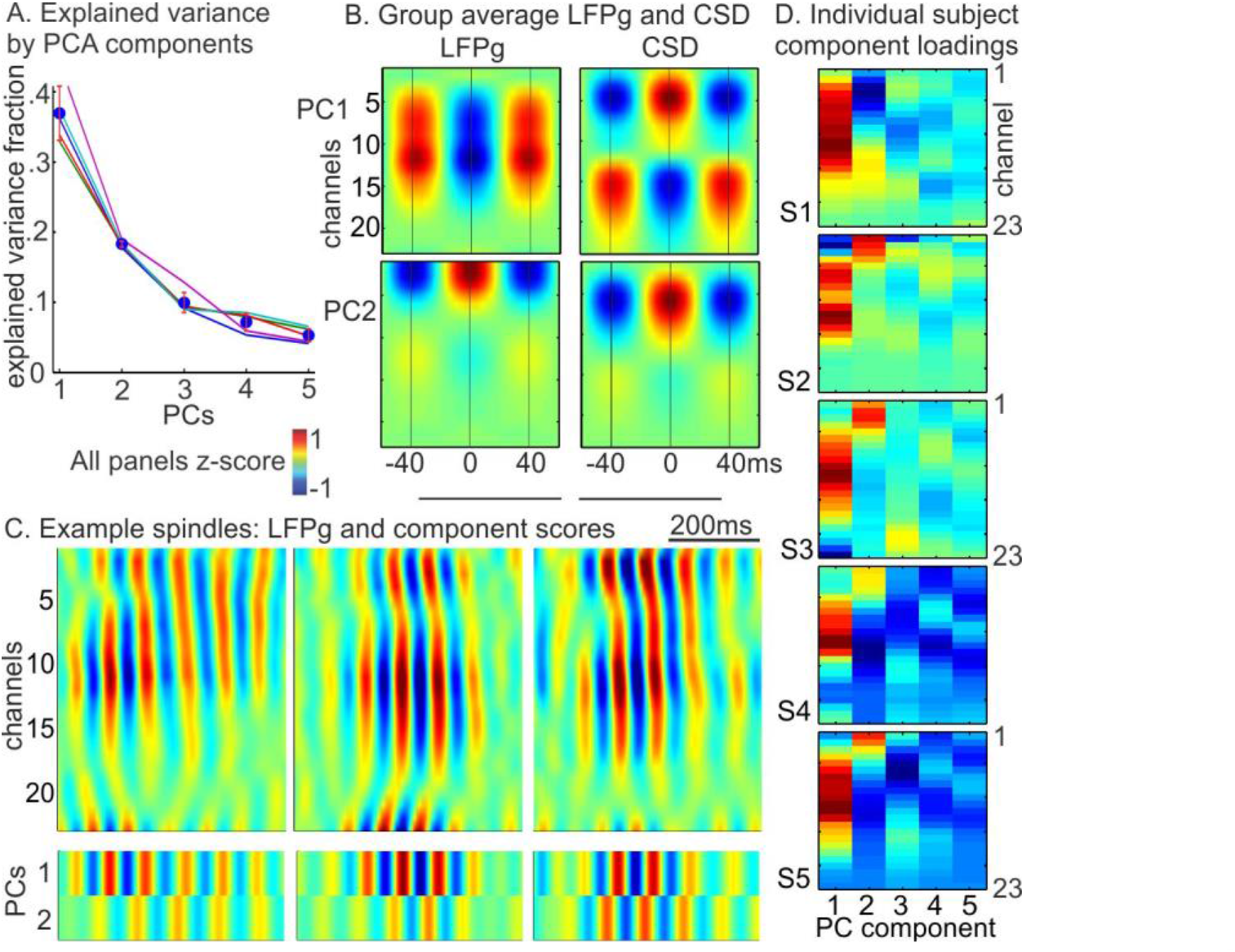
Principal component analysis. A. Fractional explained variance. Circles and error bars represent the group average and the 95% confidence interval, respectively. Lines represent the explained variance curves for each subject. B. Group average, normalized LFPg and CSD for first two principal components. Component time courses were multiplied by the channel weights then averaged across spindle epochs and then across subjects. LFPg and CSD profiles of components 1 and 2 are similar to those found for middle and upper spindles, respectively (see fig. 4). C. Example spindle waveforms (band-pass filtered, 10-16 Hz) from one subject (S3) with the corresponding time courses for PC1 and PC2 on bottom. Red indicates sinks (CSD), surface positive (LFPg) or positive PC scores. D. Individual subject component loadings show very similar channel profiles across subjects for components 1 and 2, but are weak and irregular for later components. Note that polarity (red or blue) is arbitrary so component 2 in subject 1 is similar to other subjects.

## Discussion

Intracortical laminar recordings during sleep allowed us to investigate the distribution of spindle occurrence across cortical layers. We found that spindle amplitudes and the rate of occurrence was greatest in middle channels, slightly weaker in upper channels, and greatly reduced in deep channels. Individual laminar recordings exhibited distinctive patterns of spindle co-occurrence, with spindles sometimes being detectable only in upper or only middle channels, but often occurring in both, and with spindle onset and peak latency delayed in upper channels by ∼20-50 msec on average. There were slight differences in peak spindle band frequency for upper and middle channel spindles, but each was associated with strong low-frequency power in upper channels. Factor analysis (PCA) confirmed the impression of two main consistent patterns of CSD. In all subjects, the first and second principal components corresponded to the middle and upper channel spindles, respectively. These components accounted for about half of the explained variance, with subsequent components inconsistent and accounting for little variance.

These results establish the existence of two main spindle generators. Since spindles are thought to be generated by the rhythmic activation of thalamic synapses onto cortical neurons, this implies that two thalamo-cortical systems are engaged by spindles, with differing laminar distributions of thalamocortical projections. It has previously been proposed that spindles can be generated by either or both of the two major thalamocortical systems, termed ‘core’ and ‘matrix’ (Piantoni et al., 2016) The core system is focal and acts as a relay, the matrix system is distributed and synchronizes widespread areas (Jones, 1998, Zikopoulos and Barbas, 2007). Since the core system projects mainly to middle layers and the matrix to upper, it is tempting to identify the middle channel spindles as core and the upper channel as matrix. CSD analysis provides only limited direct support for this interpretation, inasmuch as the main transmembrane current sink inferred to underlie both middle and upper channel spindle generation is indistinguishable in the group average. However, the profiles do differ strongly in the presence of a deep return current source, which was found only for the middle channel spindles. Since CSD profiles are predominantly controlled by the geometry of the receiving neuron (Einevoll et al., 2007), this strongly suggests that middle channel spindles are generated by pyramidal cells which span the layers containing the sink and source. Most cells satisfying that criterion are layer Vb pyramidal cells, whose apical dendrites extend to near the cortical surface. Furthermore, these cells have very extensive apical and basilar dendritic arborizations in the regions of the current sink and source of middle channel spindles (Ramaswamy and Markram, 2015). This configuration of large dendritic domains separated by a long narrow tube is ideal for generating source-sink pairs similar to that observed for the middle channel spindles. In contrast, upper channel spindles lack a return source in the group average, but some individual subjects may have sources above or below the sink. The presence or absence of a very superficial source is highly sensitive to how well the thin superficial layers are sampled and the exact assumptions of the CSD calculation. We noted that the superficial source was more common when the direct calculation procedure was used (Ulbert et al., 2001), rather than the inverse estimation procedure (Pettersen et al., 2006) followed here. In any case, the lack of a deep return source in upper channel spindles suggests generation by superficial pyramids. For both spindle types, high gamma modulation was greater in superficial layers, implying that phase-locking of neuronal firing is greater in those layers (Lachaux et al., 2012), as has been found in rats (Peyrache et al., 2011). Laminar gamma distribution also differed between middle and upper channel spindles. Both high and low gamma increased in upper layers during both spindle types. However, for only the middle channel sink, this increase extended to middle layers and probably the upper part of infragranular layers, providing additional evidence identifying upper channel spindles with layer II/III pyramidal cells, and middle channel spindles with layer Vb (as well as layer II/II) pyramids.

Identification of the middle and upper channel spindles with core and matrix thalamocortical afferents, respectively, is broadly consistent with the earlier studies in unanesthetized cats of (Spencer and Brookhart, 1961) and rats (Kandel, 1997 #7348} discussed in the Introduction. However, further studies are needed to confirm the identification of particular spindle patterns with thalamocortical projection systems. The laminar pattern of spindles induced by core or matrix thalamocortical input is unlikely to reflect only the layers where their respective axons terminate. A large proportion of these inputs are to interneurons, which are often not directly visible in CSD because of their dendritic configurations. This has been shown experimentally for core inputs to excitatory interneurons in layer IV (Einevoll et al., 2007), but would apply equally to matrix inputs to inhibitory interneurons in superficial layers, which include interneurons which strongly inhibit other interneurons (Cruikshank et al., 2012, Lee et al., 2015). In addition, the same voltage-gated H and T currents which underlie spindle generation in the thalamus are present in high levels in layer Vb and other cortical pyramidal cells (Ramaswamy and Markram, 2015). These and other currents result in intrinsic oscillations which can resonate with external inputs and local circuits to produce spindle-range frequencies (Silva et al., 1991). Thus, pending further clarification from experimental and modeling studies, the identification of middle and upper channel spindles with core and matrix systems must remain tentative.

Spindles have also been dichotomized into to slow vs fast, in both humans and rodents. The dichotomy in rodents is clear, with slow spindles much lower frequency (centered at ∼8 vs ∼14Hz), higher amplitude, and often epileptiform in nature (Polack and Charpier, 2006, Johnson et al., 2010). The dichotomy in humans may be less clear. Spindles are on average ∼1Hz slower over frontal cortex, in EEG, MEG (Dehghani et al., 2011b), and intracranial recordings (Andrillon et al., 2011, Piantoni et al., 2017). The individual waves in spindles also vary in frequency, with later waves also ∼1Hz slower on average (Dehghani et al., 2011b). However, using the usual 12Hz boundary, most cortical locations, and most spindle bursts, include both fast and slow spindle waves. The strongest dichotomy between slow and fast EEG spindles in humans is that slow spindles are reported to precede downstates, while fast follow (Molle et al., 2002). However, this has not been confirmed in SEEG where both fast and slow spindles clearly follow downstates (Mak-McCully et al., 2017). In our laminar recordings, upper channel spindles were slightly slower than middle, but their average frequency did not differ significantly, with both upper and middle spindles including both slow and fast spindle waves, in a continuum rather than a dichotomy. It would appear that both correspond to fast spindles in rodents, which are those associated with memory replay and consolidation (Eschenko et al., 2006, Johnson et al., 2010).

In addition to their overlap in the frequency domain, middle and upper channel spindles were closely intertwined in both location and time. Individual spindles could be mostly middle or mostly upper, but half or more had at least some involvement of both. Although on average middle spindles slightly led upper, this varied greatly across spindle epochs. Relative phase between middle and upper spindles was non-zero for most subjects but with the direction varying across subjects. Assuming that the middle channel spindles are core and the upper channel spindles are matrix, then the strong, punctate connections of the core system may support the development of oscillations within a limited thalamocortical domain, whereas the more diffuse matrix projections may have a modulatory role, perhaps spreading and synchronizing these oscillations between domains. The contribution of both core and matrix afferents in the generation of spindles would thus explain the mixture of focal and distributed spindles observed previously with intracranial recordings (Andrillon et al., 2011, Piantoni et al., 2017), and with MEG versus scalp EEG (Dehghani et al., 2010, Dehghani et al., 2011a).

This study is limited by the relatively small sample size, due to stringent clinical and technical requirements. Also, because these recordings are only performed in subjects with intractable epilepsy, close to the seizure focus, the generalizability of these results is a concern. However, we eliminated from consideration recordings with abnormal background activity, as well as epochs with epileptiform transients, or sessions following electrographic seizures. Another limitation is unavoidable ambiguities in spindle detection. We avoided false positives by excluding events that had relatively high amplitude in lower or higher frequency bands. Conversely, avoiding false negatives is important to accurately detect co-occurrence across channels. We used a relatively low initial threshold in order to detect spindles in as many channels as possible, but then excluded the spindle epochs in which the maximum amplitude in any one channel was below the median. We found that the weaker “spindles” had very low co-occurrence across channels. Given the laminar arrangement of channels, a true spindle occurring in one channel should also very likely be observed in neighboring channels, indicating that many of those weaker spindles were likely chance occurrences in single channels. Also, as noted above, although consistent phase differences between channels were observed within-individuals, generalizations regarding their direction were obscured by presumed phase-wrapping.

In conclusion, there are at least two spindle generators with differing laminar profiles of local field potentials and gamma activation. One pattern activated mainly upper layers, the other both middle and upper layers, consistent with possible generation by the core and matrix thalamo-cortical systems. Across individual spindles, the generator patterns could occur in isolation, but commonly co-occurred in various patterns. Functionally, the loose coupling of cortical spindle generators may provide a mechanism whereby spindles can integrate replay of the more focal bottom-up and distributed top-down aspects of memory (Eschenko et al., 2006, Johnson et al., 2010), leading to consolidation of coherent memories (Larkum, 2013), and global cognitive integration (Manoach et al., 2016).

## Acknowledgements

This work was supported by the National Institutes of Health Grants R01-MH-099645, R01-EB-009282, U.S. Office of Naval Research Grant N00014-13-1-0672, the MGH Executive Council on Research, Hungarian National Brain Research Program grant KTIA_13_NAP-A-IV/1-4,6, EU FP7 600925 NeuroSeeker, and Hungarian Government grants KTIA-NAP 13-1-2013-0001, OTKA PD101754. The authors thank Burke Rosen, ChunMao Wang, Adam Niese, Maxim Bazhenov, and Terrence Sejnowski for commentary, feedback and technical support.

